# Architecture and mechanism of metazoan retromer:SNX3 tubular coat assembly

**DOI:** 10.1101/2020.11.28.401588

**Authors:** Natalya Leneva, Oleksiy Kovtun, Dustin R. Morado, John A. G. Briggs, David J. Owen

**Author notes:** Correspondence should be addressed to NL, OK JAGB or DJO.

## Abstract

Retromer is a master regulator of cargo retrieval from endosomes, which is critical for many cellular processes including signalling, immunity, neuroprotection and virus infection. To function in different trafficking routes, retromer core (VPS26/VPS29/VPS35) assembles with a range of sorting nexins to generate tubular carriers and incorporate assorted cargoes. We elucidate the structural basis of membrane remodelling and coupled cargo recognition by assembling metazoan and fungal retromer core trimers on cargo-containing membranes with sorting nexin adaptor SNX3 and determining their structures using cryo-electron tomography. Assembly leads to formation of tubular carriers in the absence of canonical membrane curvature drivers. Interfaces in the retromer coat provide a structural explanation for Parkinson’s disease-linked mutations. We demonstrate that retromer core trimer forms an invariant, evolutionarily-conserved scaffold that can incorporate different auxiliary membrane adaptors by changing its mode of membrane recruitment, so modulating membrane bending and cargo incorporation and thereby allowing retromer to traffic assorted cargoes along different cellular transport routes.

---

The protein coat complex known as retromer is a central component of the endosomal sorting machinery^1,2^. Endosomal sorting is key to cellular homeostasis and its malfunction is associated with pathophysiological conditions including neurodegenerative disorders, with genetic causes being mapped to the retromer complex and its auxiliary proteins^3^. Genome-wide screens have identified retromer as one of the most important host factors for SARS-CoV-2 infection^4,5^. The core of retromer is a trimer comprising vacuolar protein sorting (VPS) proteins VPS35, VPS26, and VPS29. Cryo-electron tomography revealed the architecture of fungal retromer core assembled on membranes via a membrane adaptor, a dimer of sorting nexins (SNX) containing a Bin1/Amphiphysin/Rvs (BAR) domains, in which the core trimer forms an arch-like structure over a layer of SNX-BAR adaptors^6^. BAR domains are known for their curvature generating/stabilising properties^7^ and the accepted dogma is that membrane curvature enabling the formation of tubules is induced by the SNX-BARs, with retromer core trimer playing an auxiliary role^8^.

The core trimer has been reported to assemble with other members of the SNX-family of adaptors^9,10^ in order to expand the repertoire of cargoes that can be sorted into retromer-coated tubules and to mediate additional trafficking routes that are distinct from the retromer:SNX-BARs pathway^11–13^. In metazoa, other adaptors include SNX3, SNX12^14,15^ and SNX27^16–18^ that belong to different classes of proteins within the same SNX family. In fungi, the only known non-BAR-containing SNX adaptor is the SNX3 homologue, Grd19^13,19^. The complexity of retromer biology in metazoa likely reflects both the increased number of cargoes and increase in diversity of trafficking routes from endosomes^1,18,20^.

The current dogma contains a contradiction in that curvature generation is ascribed to the SNX-BAR adaptor proteins, but retromer’s ability to transport a wide range of cargoes has been ascribed to the use of a range of different adaptors, including those which do not contain membrane-bending BAR domains. This has led to the suggestion that metazoan and fungal retromer may adopt different architectures^21^, or that retromer has a different mechanism of action with non-BAR containing adaptors. We set out to resolve this contradiction by determining the architectures of metazoan and fungal retromer coats with non-BAR adaptor proteins.

## The structure of the retromer:SNX3 membrane coat

We expressed and purified recombinant metazoan and fungal retromer core trimer, as well as the membrane adaptors, metazoan SNX3 and its fungal homologue Grd19. SNX3/Grd19 contains no BAR domain, consisting only of a phox-homology (PX) domain that selectively binds phosphatidylinositol 3-phosphate (PI(3)*P*)^22^. Each core trimer was reconstituted with its respective adaptor on membranes containing the early endosomal marker phospholipid PI(3)*P*^23^ and cargo peptides (**Fig S1a**) derived from the C-terminal domains of Wntless (Wls) for metazoa^11,24,25^ or the Ca^2+^-dependent serine protease Kex2^26–28^ for fungi. Cryo-electron microscopy imaging of these membrane-reconstituted complexes revealed the formation of abundant long tubules with a dense protein coat (**Fig S1b**) immediately demonstrating that SNX-BARs are not required for tubulation of the underlying membrane.

We imaged the protein-coated tubules by cryo-electron tomography (cryo-ET) and applied reference-free subtomogram averaging (STA) to generate electron microscopy (EM) density maps of the assembled coat. Initial alignments showed the presence of arch-like units similar to those previously observed in fungal retromer:SNX-BAR coats and on protein-coated tubules within green algae^6^. Using local alignment we produced overlapping maps (**Fig S2**) that revealed details of the fully assembled cargo-containing coats of metazoan retromer:SNX3 and fungal retromer:Grd19 at subnanometer resolution (**Fig 1**). Consistent with the measured resolution (**Fig S2**), protein secondary structure elements were readily resolved in the EM maps allowing unambiguous docking of experimental and homology atomic models. Comparison of the assemblies demonstrated that metazoan (**Fig 1b**) and fungal (**Fig 1d**) coats are essentially identical and hereafter we describe metazoan retromer:SNX3 unless stated otherwise. The coat consists of arch-like units formed by VPS35 homo-dimerisation that are directly connected to membrane-attached assemblies of VPS26 dimers and two SNX3 molecules (**Fig 1b,d**) contradicting recent assertions that the VPS35 arch is incompatible with SNX3 membrane binding and that retromer:SNX3 must therefore adopt a flat structure^21^. Arches were arranged in a pseudo-helical lattice wound around membrane tubules (**Fig 1a,c**).

**Fig 1.**
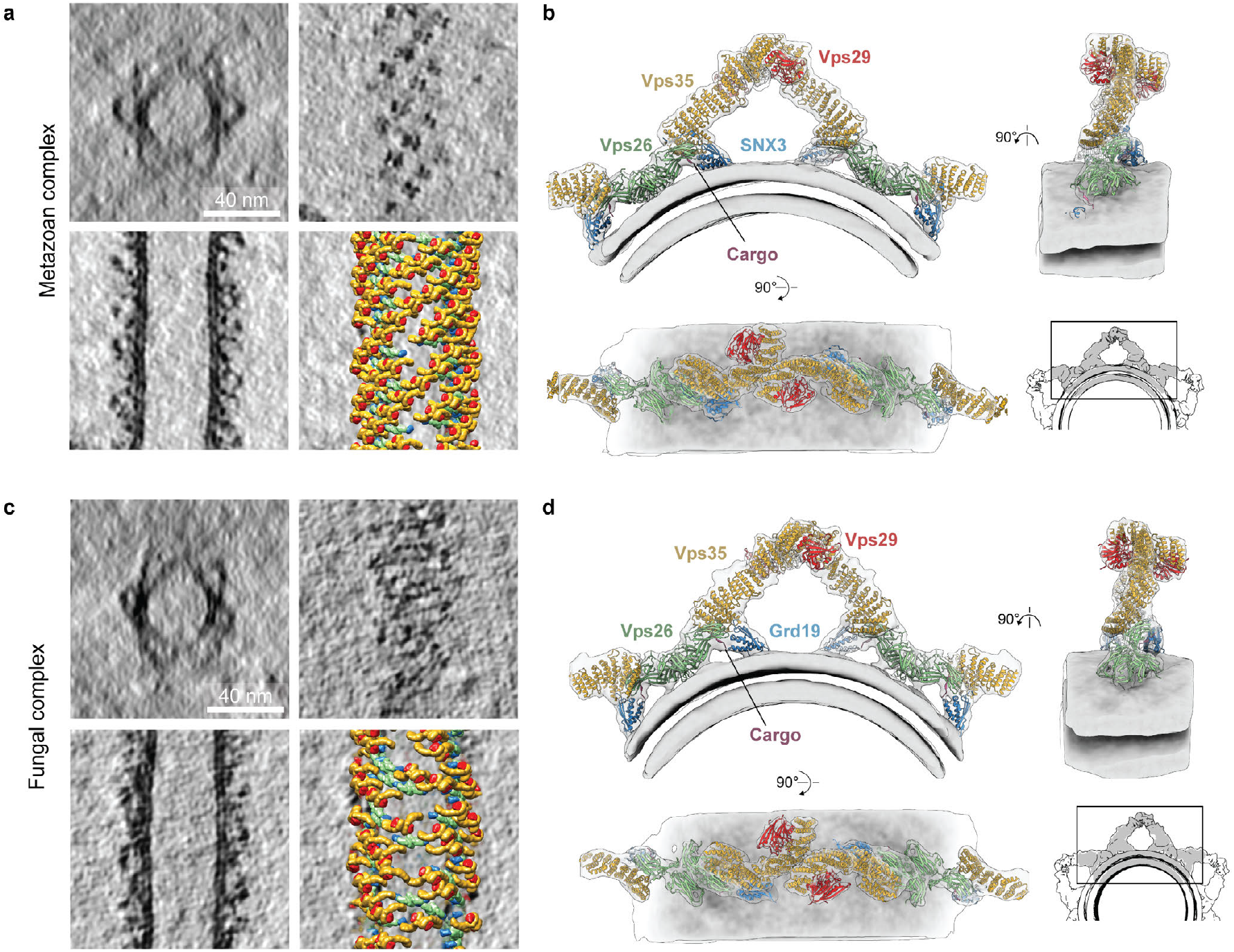
Overview of metazoan and fungal retromer:SNX3 coats assembled on membranes in the presence of cargo peptides. **a,** Representative slices through tomographic reconstructions of metazoan retromer:SNX3-coated membrane tubules. Upper left panel: radial cross-section through a tubule. Upper right: axial cross section at the level of arch heads. Lower left: axial cross-section through the middle of a tubule. Lower right: the slice from the lower left overlaid with models of coat subunits placed in space according to their positions and orientations found thorough STA. Models colored as in **b**. **b**, Overlay of ribbon-depicted atomic models colour-coded by protein, and composite EM map covering the membrane, arch and two neighboring VPS26 dimer regions with short segments of VPS35 of the next arches. The grayscale inset shows a model of a section of a tube prepared by overlay of three composite maps. **c,d,** as in **a,b** for the fungal retromer:Grd19 complex.

## Membrane interactions of the retromer:SNX3 coat

In assembled retromer:SNX-BAR, the core trimer does not contact the membrane but is bound to the SNX-BAR adaptor layer via loops 5 and 9 of VPS26^6^. In contrast, the core trimer in retromer:SNX3 coats docks directly to the membrane and we now see the structural details of this docking (**Fig 2**): loops 5, 6, 9 and 15, and the N-terminus of VPS26 directly contact the membrane (**Fig 2b, Fig S3a**). Phosphorylation in VPS26 loop 6 was previously shown to influence the trafficking of chitin synthase 3 in yeast^29^. Structurally, VPS26 loops 6 and 15 are equivalent to the β-arrestin ‘finger’ and ‘C-loop’ which interact directly with the transmembrane core and membrane proximal residues of cargo receptor^30^. In the context of retromer:SNX3 coat, loops 6 and 15 of VPS26 have access to the membrane and we speculate that they play a similar role interacting with transmembrane domains of cargoes as described for TGN38 in metazoa^31^ and Snc1 in yeast^32^.

**Fig 2.**
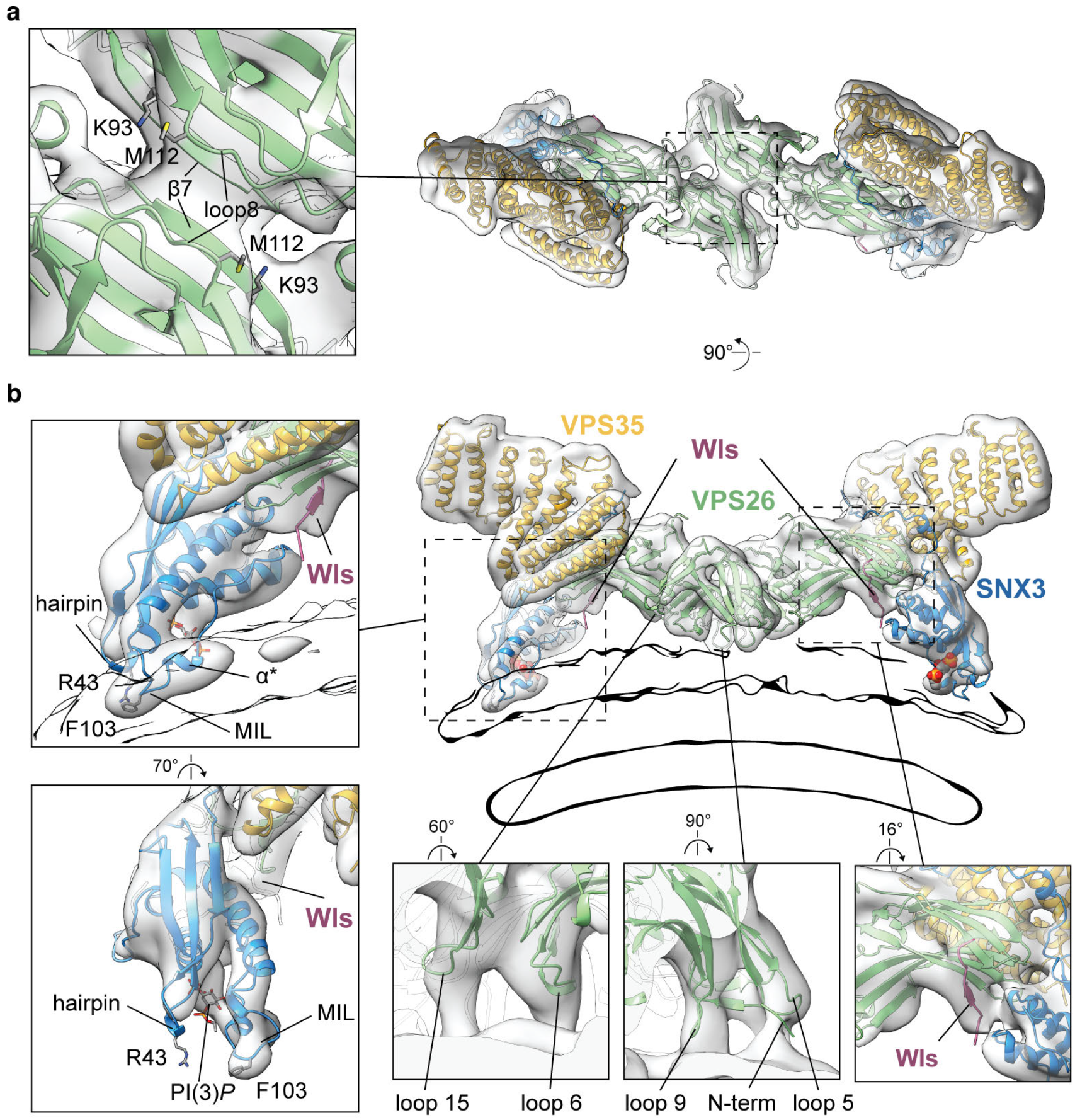
Structure of the membrane-proximal region of the metazoan retromer:SNX3 complex. Ribbon models colored by subunit and fitted within semi-transparent EM density maps. Modelled PI(3)*P* is shown as spheres (in overviews) or stick model (in close up views). **a**, View looking ‘down’ onto the membrane. Inset shows a close-up of the VPS26 homo-dimerisation interface, marking secondary structure elements forming the interface and residues with PD-associated mutants. **b**, View perpendicular to membrane. Inset panels show regions indicated by boxes. The membrane is shown as an outline of the same EM map at lower contour level. Two left hand panels illustrate features involved in SNX3 membrane anchoring including the membrane-submerged MIL, α*, and bound PI(3)*P*. The conserved R43 and F103 within the MIL are marked. Lower panels demonstrate membrane-contacting VPS26 loops (left, middle panels shown at lower contour level than the overview) and EM density for the Wls-occupied cargo-binding site (right panel). Equivalent views for the fungal retromer:Grd19 complex are shown in **Fig. S3b**.

To engage with sorting motifs located on the unstructured cytoplasmic tail of cargo, VPS26 adopts an ‘open’ form where the outward movement of the β10 strand generates the interface for cargo binding^33^ as opposed to the closed conformation observed in the absence of cargo^34^ (**Fig S4**). Previously, this interface was described by crystallizing a minimal part of the retromer complex with a cargo motif covalently linked to the VPS26 as a chimaera^33^. For both metazoan and fungal retromer:SNX3 coat reconstitutions we included membrane-attached cargo peptides that contain a canonical *Øx(L/M)* sorting motif (where *Ø* is an aromatic amino acid and *x* any residue). In both structures, we observed that VPS26 is in the open conformation (**Fig S4**) and that density corresponding to the cargo peptide (Wls and Kex2, respectively) is visible on VPS26 near the SNX3 binding interface (**Fig 2b, Fig S3b**), indicating that this site can bind cytoplasmic portions of cargoes in a physiologically relevant, membrane-assembled state.

VPS26 forms a homo-dimer on the membrane by β-sheet complementation between the β7 strands of the N-subdomains creating an interface identical to that seen in the retromer:SNX-BAR coat^6^. VPS26 homo-dimerisation is therefore not specific to interaction with a SNX-BAR layer as has been suggested^21^. Loop 8, which is positioned above β7, is also likely involved in the stabilisation of VPS26 dimers (**Fig 2a**). Several mutations in VPS26 (K93E, M112V/M112I) occur in sporadic and atypical Parkinson’s disease (PD) with an unknown causative mechanism^35^. The coat assemblies described here place these mutations in close proximity to the VPS26 homo-dimerisation interface (**Fig 2a**) where they will likely impair VPS26 dimerisation and hence coat formation providing an explanation for the mutations’ pathophysiological effects.

In the structure presented here, SNX3 binds the core trimer at the interface between VPS26 and VPS35. It is oriented with the PI(3)*P* binding pocket facing the membrane and with a clear density for the PI(3)*P* head group present in the pocket (**Fig 2b**). SNX3 is further anchored in the membrane via the <u>M</u>embrane <u>I</u>nsertion <u>L</u>oop (MIL) containing the short α-helix α*^36^, and the β1–β2 hairpin, which contains a membrane-facing arginine (**Fig 2b, Fig S3a**). The fungal Grd19 is similarly anchored, although it has a shorter MIL compared to SNX3 and lacks α* (**Fig S3b,c**). A sequence alignment of PX domains from all known SNX-family adaptors confirms that although variable in length, the MIL is a consistent feature of membrane interacting PX domains (**Fig S3c,d**). In contrast, in PX domains of SNX5 and SNX6 that are unable to bind membranes^22^, the MIL is replaced by an extended helix-turn-helix structure involved in cargo binding (**Fig S3c,d**). In order to perform their function in cargo binding and sorting, SNX5 and SNX6 form heterodimers with either SNX1 or SNX2^37^, both of which have MIL-containing PX domains.

## The retromer arch is asymmetric and conserved

Using local reconstruction and 3D classification, we found that for both metazoan and fungal retromer:SNX3 datasets the arch is asymmetrical (**Fig 3**). One VPS35 monomer is more curved that the other and the two monomers interact via an asymmetric VPS35:VPS35 dimerisation interface. This asymmetry causes the arches to tilt by approximately 22° away from the perpendicular relative to the membrane (**Fig 1b,d**). We re-processed the retromer:SNX-BAR dataset (**Fig S2a)** where we had previously applied 2-fold symmetry and found that it, in fact, also uses the same asymmetric VPS35 dimerisation interface (**Fig S5**). The interface is formed predominantly by the loops between α28-α29, α30-α31 and α33-α34 from the straighter conformation of VPS35 binding to α30 and α28 of the more curved VPS35 molecule (**Fig 3, Fig S5**). The interface relies on electrostatic and hydrophobic contacts between highly conserved residues with a prominent electrostatic bridge formed between E615, D616 and E617 on straighter VPS35 and K659 and K663 on the more curved VPS35 (**Fig 3b,c, Fig S6c,e**). The combination of those residues was confirmed biochemically to be crucial for the formation of dimerisation interfaces^21^, however their proposed model of dimerisation was different (**Fig S6d,e**). Intriguingly, VPS35L, which is predicted to be the VPS35 homologue in the Retriever complex^1,38^ does not contain the critical dimerization residues (**Fig S6e**), suggesting that Retriever might not homodimerize in the same manner and may be unable to form arches.

**Fig 3.**
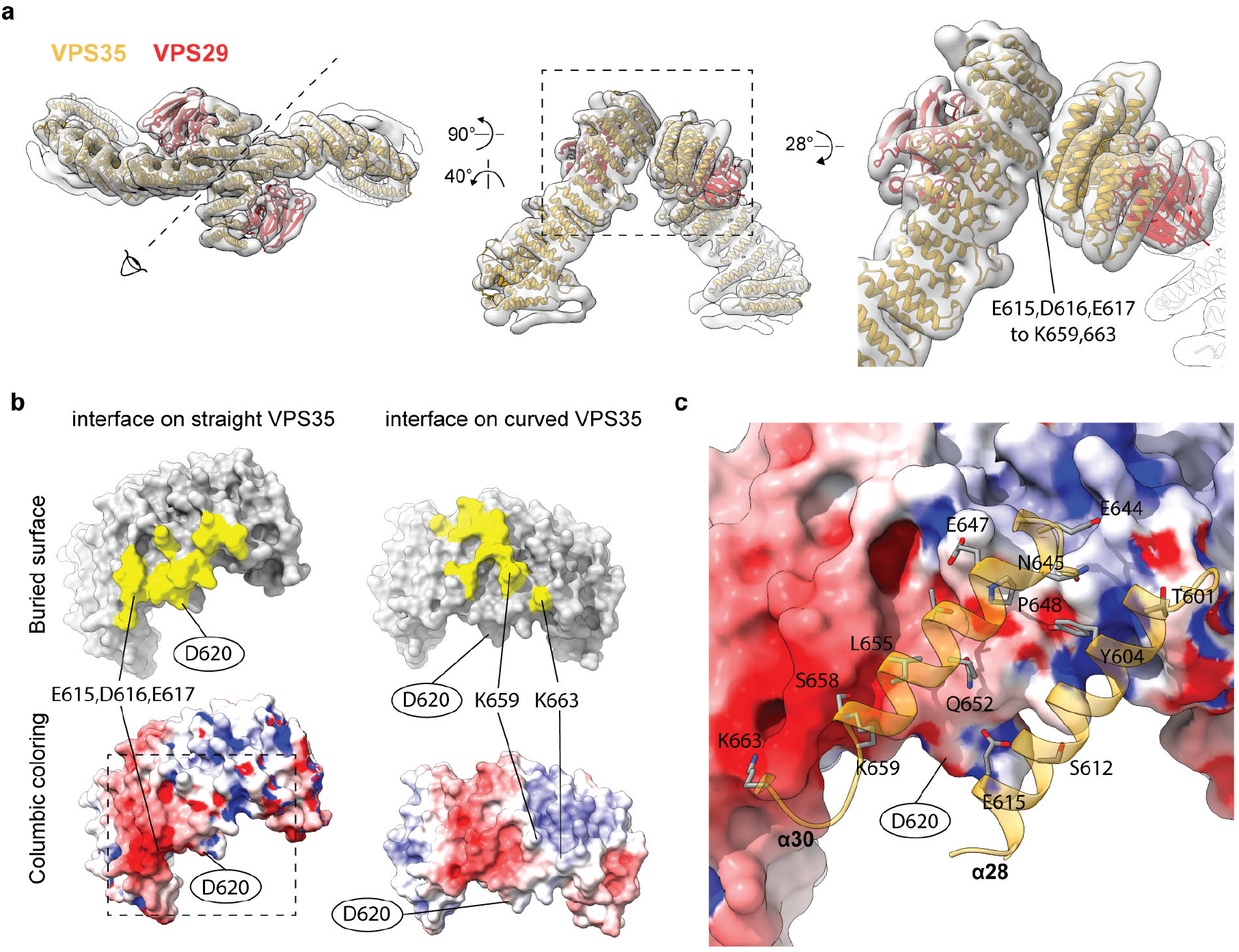
Conserved, asymmetric VPS35 dimerisation interface. **a,** Top and side views of ribbon representation of flexibly fitted VPS35/VPS29 colour-coded models into the EM map of the metazoan arch. The direction of view presented in the middle panel is indicated in the left panel. The right hand panel shows a close-up of the area boxed in the middle panel. The EM density corresponding to the electrostatic interaction between E615, D616, and E617, and K659 and K663 is indicated. **b**, Cutaway views of the dimerisation surfaces of straighter and the more curved VPS35 molecules. Model-generated surfaces are colored by the buried interface in yellow (top panels) or by columbic potential from red (negative) to blue (positive) (bottom panels). Key electrostatic residues and D620 which is mutated in PD (oval outline) are indicated. To produce the interface model preserving experimentally determined side chains orientations, the atomic model of the C-terminal region of human VPS35 (PDB 2R17:B^47^) was docked as a rigid body into the EM densities of the helices facing the interface of both VPS35 molecules. **c**, Close-up view of the surface model of the interface on the straight VPS35 (boxed area in **b**) overlaid with ribbon model of key contact helices α28 and α30 in the curved VPS35 with a stick representation of residues within the buried interface.

Asymmetric VPS35 dimerisation has possible implications for retromer function. It leads to an asymmetric VPS35 surface at the apex of the arch to which cofactors could bind directionally and with a stoichiometry of 1 cofactor: 2 VPS35 molecules. For example, a D620N mutation that is associated with familial autosomal dominant and sporadic PD^39^ causes loss of affinity to the Family with sequence similarity 21 (FAM21) protein^40^, the retromer-binding subunit of Wiskott-Aldrich Syndrome Protein and SCAR Homolog (WASH) complex^41^. D620 is part of the buried interface in straighter VPS35, but is exposed in the more curved VPS35 (**Fig 3b,c**), suggesting that there is only one FAM21 binding site at the apex. The interfaces of the two VPS29 subunits at the apex of the arch may also be differently accessible for binding by regulator factors such as the Tre-2/Bub2/Cdc16 domain family member 5 (TBC1D5)^42^, Vps9-ankyrin repeat protein (VARP)^43^ and RidL, the effector of pathogenic bacteria *Legionella pneumophila*^44^. Asymmetry could impose polarity and therefore directionality of coat assembly.

## The retromer coat is a modular assembly of a conserved scaffold and alternative adaptors

The recent observation that retromer forms lower-order oligomers on bilayers with limited flexibility^45^ suggests that higher-order oligomerization and membrane remodelling are interconnected processes. We observed that on flexible membranes, retromer forms tall arches and not the recently suggested flat structures (**Fig. S6a,b**). Arches are assembled into linear chains via two homo-dimerization interfaces - the asymmetrical VPS35 dimer interface and the symmetrical VPS26 dimer interface (**Fig 1**). This mechanism of coat formation is invariant in the retromer coats we have studied, independent of kingdom (animals, fungi and plants), and independent of adaptor (with or without BAR domain) (**Fig 4, Fig S7**). This invariable arched scaffold can employ two distinct membrane coupling modes to accommodate different adaptor types: it can bind either directly to the membrane (retromer:PX) or via the BAR layer (retromer:PX-BAR), **Fig 4a.** The direct membrane-binding mode can be observed for retromer arches within cells (**Fig S7**). We can now directly compare coats with different membrane binding modes to understand the mechanism of the membrane tubulation.

**Fig 4.**
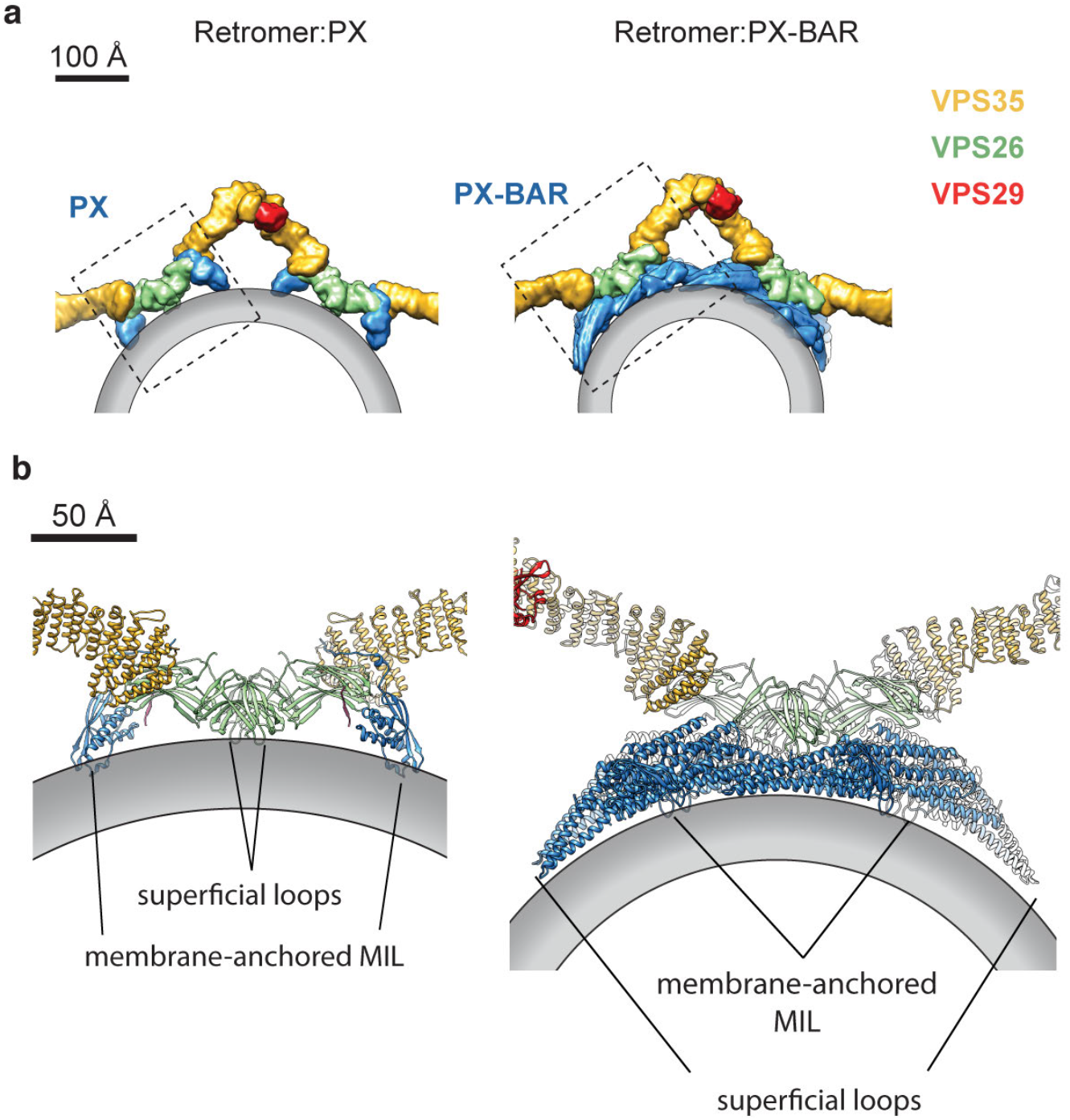
Modularity of the retromer coat allows distinct modes of membrane attachment using different adaptors. **a**, Surface model illustration of retromer:PX (modeled on metazoan retromer:SNX3) and retromer:PX-BAR (modeled on fungal retromer:Vps5). Surface models were generated from pseudoatomic models of corresponding complexes and color-coded. The membrane bilayer is schematically shown as a grey outline. **b**, Close-ups of the areas boxed in **a** showing the assembly of membrane-attached subunits in each complex. The VPS26/SNX3 assembly (left) and the four Vps5 dimers contacted by the VPS26 dimer (PDB 6W7H^6^) (right) have inherently curved structures that use a combination of membrane-anchored MILs (from the PX domains of SNX3 and Vps5) and superficially-attached loops (from VPS26 and the tip-loops of the BAR domain) to contact the membrane.

In the case of retromer:SNX3 coats, membrane bending by the core trimer arches is enhanced by further scaffolding through the curved VPS26-PX (**Fig 4b**) domain surface, and by membrane insertions from VPS26 and SNX3, thus explaining the previous observation that retromer promotes the membrane remodelling activity of yeast Grd19^46^. In retromer:PX-BAR coats, membrane bending will be enhanced through the membrane re-modelling action of BAR domains (**Fig 4b**). Our model also permits retromer assemblies with other non-BAR SNX adaptors such as SNX12 and SNX27 that may have similar membrane remodelling properties to SNX3. This suggests a general modular scheme for retromer function in which the core trimer arch – the conserved architectural element of the different retromer coats – forms a scaffold that contributes significantly to membrane bending and therefore tubule formation, while additionally helping to propagate membrane curvature over long distances by oligomerisation. The core trimer can make use of a range of different adaptor modules, which determine the cargo to be incorporated and that simultaneously modulate the degree of membrane bending appropriate to the size of the cargo and/or the bilayer properties along the trafficking route.

## Materials and methods

### Protein expression and purification

All proteins were expressed at 20°C in *Escherichia coli* BL21(DE3) following induction with 0.2 mM isopropyl β-D-1-thiogalactopyranoside as previously described in^6^. Glutathione S-transferase (GST)–tagged proteins were constructed in pGEX-4T2 by genetically fusing fungal (*Chaetomium thermophilum* – Ct) CtGrd19 (Uniprot G0S0X3), mouse SNX3 (Uniprot O70492) and zebrafish VPS26 (Uniprot Q6TNP8) to the C-terminus of GST using PCR-based cloning method^48^. It allowed us to introduce an additional cleavage site for the PreScission protease located downstream of the thrombin site. Fungal core trimer was purified as described in^6^. The mouse GST-VPS35 (Uniprot Q9EQH3):VPS29 (Uniprot Q9QZ88) heterodimer was expressed and purified as in^49^. All GST-containing proteins we purified following the standard protocol^50^. In short, cells were lysed by high-pressure homogenization in 50 mM Tris-HCl pH 8.0, 200 mM NaCl buffer. The homogenate was cleared by centrifugation at 30,000g and loaded onto a gravity-flow column containing Glutathione Sepharose 4B (GE Healthcare). The proteins were eluted by protease cleavage of the GST-tag (thrombin or PreScission (Sigma-Aldrich) were used depending on the enzyme availability) and were further purified by gel-filtration chromatography on Superdex 200 10/300 column (GE Healthcare) in Buffer A (20 mM HEPES-KOH pH7.5, 200 mM NaCl, 1mM tris (2-carboxyethyl)phosphine (TCEP).

His-tagged cargoes were constructed in pRSFDuet-1 by genetically fusing residues 733-846 of Kex2 (Uniprot G0SHU5), and residues 493-541 of Wls (Uniprot Q6DID7) to the fusion-tag, resulting in His10Kex2 and His6Wls respectively. The cargo tail peptides were isolated on gravity-flow column containing Ni-NTA agarose (GE Healthcare) followed by gel-filtration chromatography on Superdex Peptide 10/300 (GE Healthcare) in Buffer A.

### Liposomes and tubulation reactions

Liposomes composed of 1,2-dioleoyl-sn-glycero-3-phosphocholine (DOPC), 1,2-dioleoyl-sn-glycero-3-phosphoethanolamine (DOPE), 1,2-dioleoyl-sn-glycero-3-phospho-L-serine (DOPS), and 1,2-dioleoyl-sn-glycero-3-[(N-(5-amino-1carboxypentyl)iminodiacetic acid)succinyl] nickel salt (DGS-NTA(Ni) (all Avanti Polar Lipids) in a 42:42:10:3 molar ratio, with 3 mol% of dipalmitoyl-phosphatidylinositol-3-phosphate (PI(3)*P*) (Echelon Biosciences) were prepared at a lipid concentration of 0.5 mg/ml in Buffer A by extrusion through a 0.4 μm polycarbonate filter (Avanti Polar Lipids). For the tubulation reaction, 2.5 μM of the core retromer trimer and 3.5 X molar excess of the adaptor (SNX3 or Grd19), and the corresponding cargo peptide (Wls or Kex2) were incubated with 0.16 mg/mL of liposomes for 4 hours at 22 °C in Buffer A.

### Cryo-electron tomography sample preparation and data acquisition

10 nm gold fiducial markers (BBI Solutions) in buffer A were added to the tubulation reaction (1:10 fiducials:reaction volume ratio). 4 μl of this mixture was back side blotted for 6 s at relative humidity 98% and temperature 19°C on a glow-discharged holey carbon grid (CF-2/1-3C, Protochips) before plunge-freezing in liquid ethane (Leica EM GP2 automatic plunger). Dose-symmetrical tilt series acquisition ^51^ was performed on an FEI Titan Krios electron microscope operated at 300 kV using a Gatan Quantum energy filter with a slit width of 20 eV and a K2 or K3 direct detector operated in counting mode. The total exposure of ~130 e^−^/Å^2^ was equally distributed between 41 tilts. 10 frame movies were acquired for each tilt. The details of data collection are given in **Table S1.** The selection of acquisition areas was guided by suitability for high resolution tomographic data collection (i.e. vitreous ice quality, lack of crystalline ice contaminations, intactness of the carbon support) and was not based on the morphology of tubules.

### Tomogram reconstruction

Image pre-processing and tomogram reconstruction were performed essentially as described in^52^. The IMOD v. 4.10.3 package^53^ was used to align frames in raw movies and correct for detector gain and pixel defects. Several tilt series in each dataset were discarded at this point (dataset sizes and all exclusions are listed in **Table S1)** due to tracking errors or large beam-induced sample movements. In addition, defective high-tilt images (due to tracking error, large objects like a grid bar, or contaminations coming in the field of view) were also removed prior to low-pass filtering to the cumulative dose^54^. Tilt series were aligned based on fiducial markers in the IMOD package. The aligned tilt series were binned four times and reconstructed by weighted back projection in IMOD resulting in the non-CTF corrected tomograms that were used for visual inspection of the quality of tomographic reconstruction, tubule picking, and defocus estimation using CTFPLOTTER (within the IMOD). To reconstruct 3D CTF-corrected tomograms for subtomogram averaging, dose filtered tilt series were CTF-corrected by phase-flipping and back projected into tomographic reconstructions using novaCTF^55^ with a 15 nm strip width. The resulting unbinned tomograms were binned by factors of two (generating bin2, bin4, bin8 tomograms) with anti-aliasing using IMOD.

### Subtomogram alignment

Subtomogram alignment and averaging were done as previously described in^6^ using MATLAB (MathWorks) functions adapted from the TOM^56^, AV3^57^. As in^52^ we used a modified wedge mask representing the amplitudes of the determined CTF and applied exposure filters at each tilt^58,59^. **Table S2** contains a summary of data processing parameters. Subtomogram averaging was performed identically and independently for the two datasets as described in the following sections.

### Extraction of initial subtomograms

Centers of coated tubules were manually traced in bin4 tomograms and their diameters recorded using Chimera (v1.14) and the custom plug-in^60^. All tubules were included (no selection was made for diameter or morphology) (**Table S1)**. The positions of subtomograms were defined on the surface of tubes with uniform sampling at every 44 Å. Initial subtomogram orientations were set to be normal to the membrane surface and with the in-plane angle perpendicular to the main tubule axis.

### *Ab initio* reference generation

Subtomograms were extracted at the initial positions from bin8 tomograms and averaged according to their initial orientations producing an initial reference. Subtomograms were then aligned to this reference in the direction perpendicular to the membrane and averaged to generate a reference containing density layers corresponding to the lipid bilayer and protein layer. An arbitrary subset of the data (10 tomograms) was then further iteratively aligned allowing both shifts and angular search. A cylindrical mask was applied passing the protein layer and the membrane. Five iterations of such alignment were performed with an 8° to 4° angular search increment and a 40 Å low-pass filter. The resulting average was shifted and rotated to place the arch or the VPS26 dimer in the centre of the box.

### Subtomogram alignment and classification

The resulting references were used to align the complete dataset with similar alignment parameters. Overlapping subtomograms resulting from oversampling at the initial extraction stage were removed by selecting the subtomogram with the highest cross-correlation score within a distance threshold of 55 Å. Subtomograms were then split by tubule into odd and even half datasets for further processing. Subsequent alignments were performed independently on the odd and even half-sets. The search space and angular increments were gradually decreased, and the low-pass filter was gradually moved towards higher resolution. After visual examination, misaligned subvolumes (those not aligned to the membrane) were removed using a cross-correlation cut-off that was manually selected for the whole dataset. In some cases, subvolumes located close to the edge of tomograms were also removed. At the end of each iteration, subtomograms within each half-set were averaged and resolution was assessed by Fourier shell correlation (FSC).

3D classification was performed using principle component analysis of wedge-masked difference maps^61^ with calculations implemented in MATLAB using code adapted from TOM^57^, AV3^62^.

### EM map postprocessing

Final half-maps were filtered with soft masks to remove the box edge using IMOD (4.10.3) and EMAN (2.2.2) packages. Local resolution was measured using relion_postprocess from the Relion 3.0.8 package^63^. The final sharpened maps were prepared using local resolution filtering and denoising implemented in LAFTER (v1.1)^64^.

### Model building

Rigid body fitting was performed in Chimera, peptide linkers were build using Modeler (v 9.24), and flexible fitting used the Real-Space Refine procedure in Phenix (1.18.2-3874). SWISS-MODEL^65^ web service was used for homology modelling. Chimera and ChimeraX (v1.0) packages were used for molecular visualisation.

To build models of the arch, structures of human VPS29 (PDB 2R17:A) and N- and C-terminal portions of VPS35 (PDB 5F0L:A and 2R17:B respectively) were fitted into the EM map of the metazoan arch region as rigid bodies. The missing linker between the two portions of VPS35 was built and the resulting full-length VPS35 model was flexibly fit with secondary structure constraints to account for movements of individual helixes occurring along the length of α-solenoid. The fungal model was built using the identical procedure, but based on a homology model of CtVPS35 (Uniprot G0S709) generated from the above human VPS35 models together with the experimental model of CtVps29 (PDB 5W8M).

To build models of the VPS26 dimer region, two copies of the experimental model of human VPS26/SNX3/N-termVPS35/DMT-II cargo peptide assembly (PDB 5F0L) were rigidly fitted into the metazoan EM map. They were then split into VPS26/DMT-II, SNX3 and N-terminal VPS35 subunits, and the N-terminal VPS35 subunit was flexibly fitted into the arch EM map as described above. Subunit positions were refined by rigid body fitting using sequential fit command in Chimera. The PI(3)*P* model was copied into SNX3 from the yeast homologue Grd19p (PDB 1OCU) complexed with PI(3)*P*. Modelling into the fungal EM map was done identically but the human subunits (excluding the DMT-II cargo peptide) were replaced with the corresponding *Ct* homology models. CtGrd19 and CtVps26 were modelled on yeast 1OCU and human 5F0L PDB models respectively.

## Acknowledgements

This study made use of electron microscopes at the MRC LMB EM Facility and the high-performance computing resources at LMB. We acknowledge Jake Grimmett and Toby Darling from MRC LMB Scientific Computing for providing technical support. We thank Kun Qu for providing assistance in data collection and advice during subtomogram data processing. NL acknowledges Brett Collins for the gift of the cDNA for CtGrd19 and zVPS26 and the Cryo-EM Facility, University of Cambridge, Department of Biochemistry and Dimitri Y. Chirgadze and Steven W. Hardwick in particular, for training and assistance during initial sample screening. The Cryo-EM Facility, Department of Biochemistry, University of Cambridge is funded by the WT 206171/Z/17/Z and 202905/Z/16/Z. NL and DJO were supported by WT grant 207455/Z/17/Z. Work in JAGB’s lab was funded by the Medical Research Council (MC_UP_1201/16) and the European Research Council (ERC) under the European Union’s Horizon 2020 research and innovation programme (ERC-CoG-648432 MEMBRANEFUSION).

## Author contributions

NL, OK, JAGB and DJO designed the project. NL performed biochemical characterisation, protein preparation and in vitro assembly reactions. NL performed subtomogram averaging with assistance from OK and JAGB. OK performed model building. DRM assisted OK and NL during data collection and data processing. NL, OK, JAGB and DJO prepared the manuscript with input from DRM.

## Extended data figures and tables

**Fig S1.**
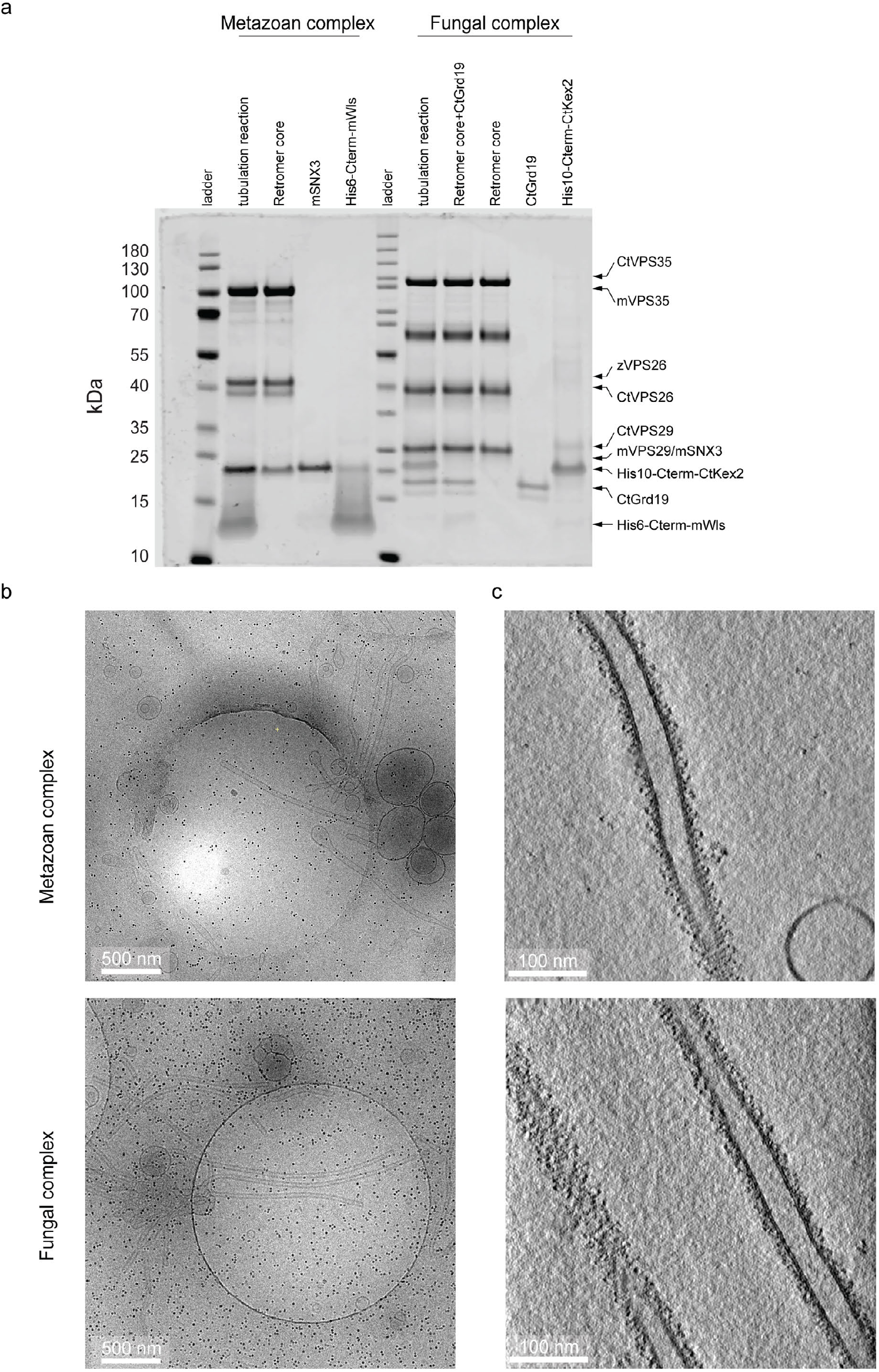
Reconstitution of retromer-coated tubules. **a.** Coomassie stained 4-12% Bolt™ Bis-Tris polyacrylamide gel analyses of individual components and of the corresponding assembled tubulation reactions. PageRuler™ Prestained Protein Ladder (left lane, cat no #26616) and PageRuler™ Unstained Protein Ladder (center lane, cat. no. #26614) were used as molecular weight rulers. Prefixes indicate protein origins: m (mouse), z (zebra fish), Ct (*Chaetomium thermophilum).* **b.** Representative cryo-EM images of reconstitution reactions for metazoan and fungal coats, and **c.** slices through representative tomograms of the respective reconstitutions.

**Fig S2.**
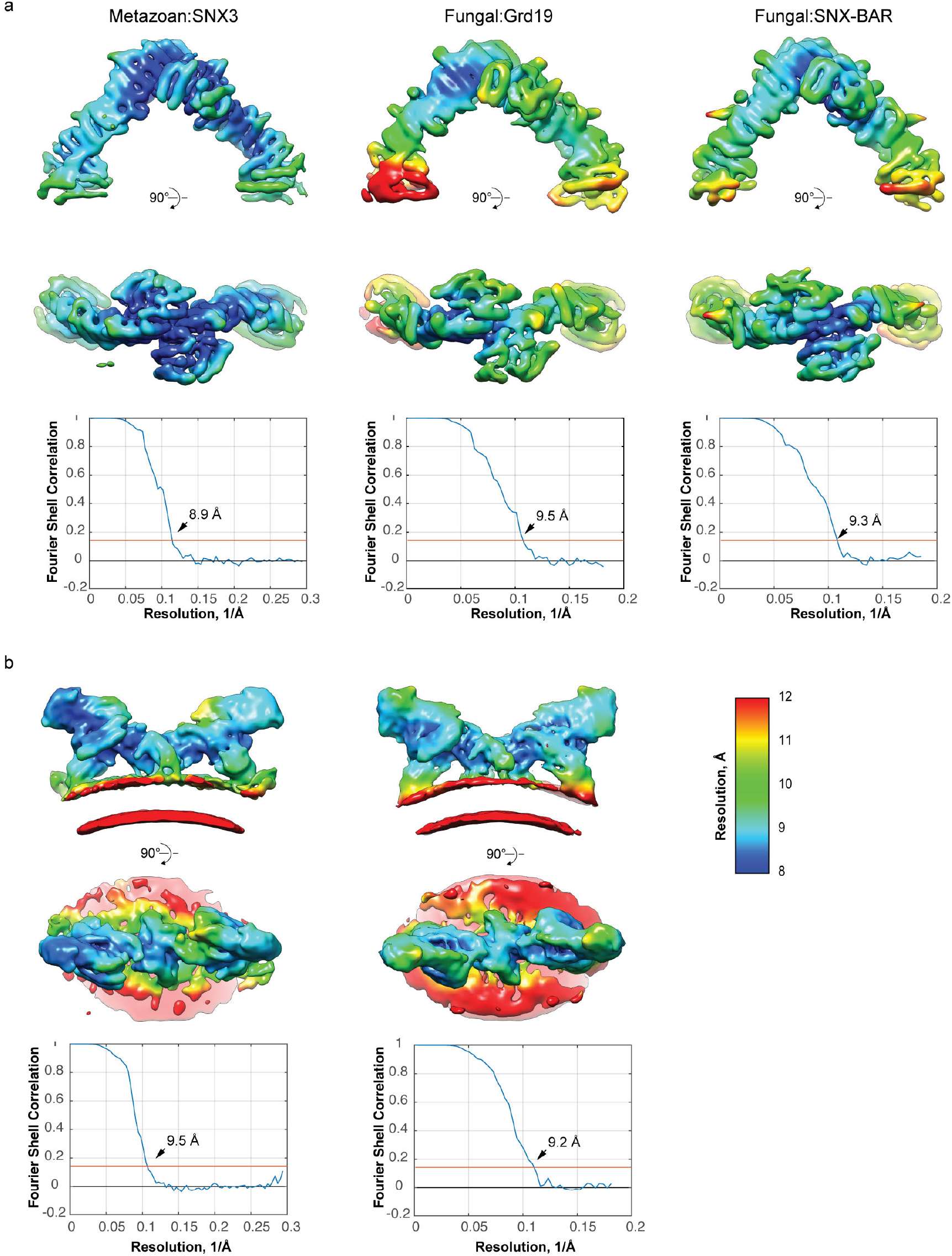
Global and local resolution of focused EM maps. Maps of arch (**a**) and VPS26 dimer (**b**) regions are colored by local resolution with corresponding mask-corrected FSC curves shown beneath each map. Metazoan:SNX3 and fungal:Grd19 map were generated from the data in this work while the map of fungal:SNX-BAR was generated by a reprocessing previously published cryo-ET data on Ct retromer:Vps5 structure (Kovtun et al 2018).

**Fig S3.**
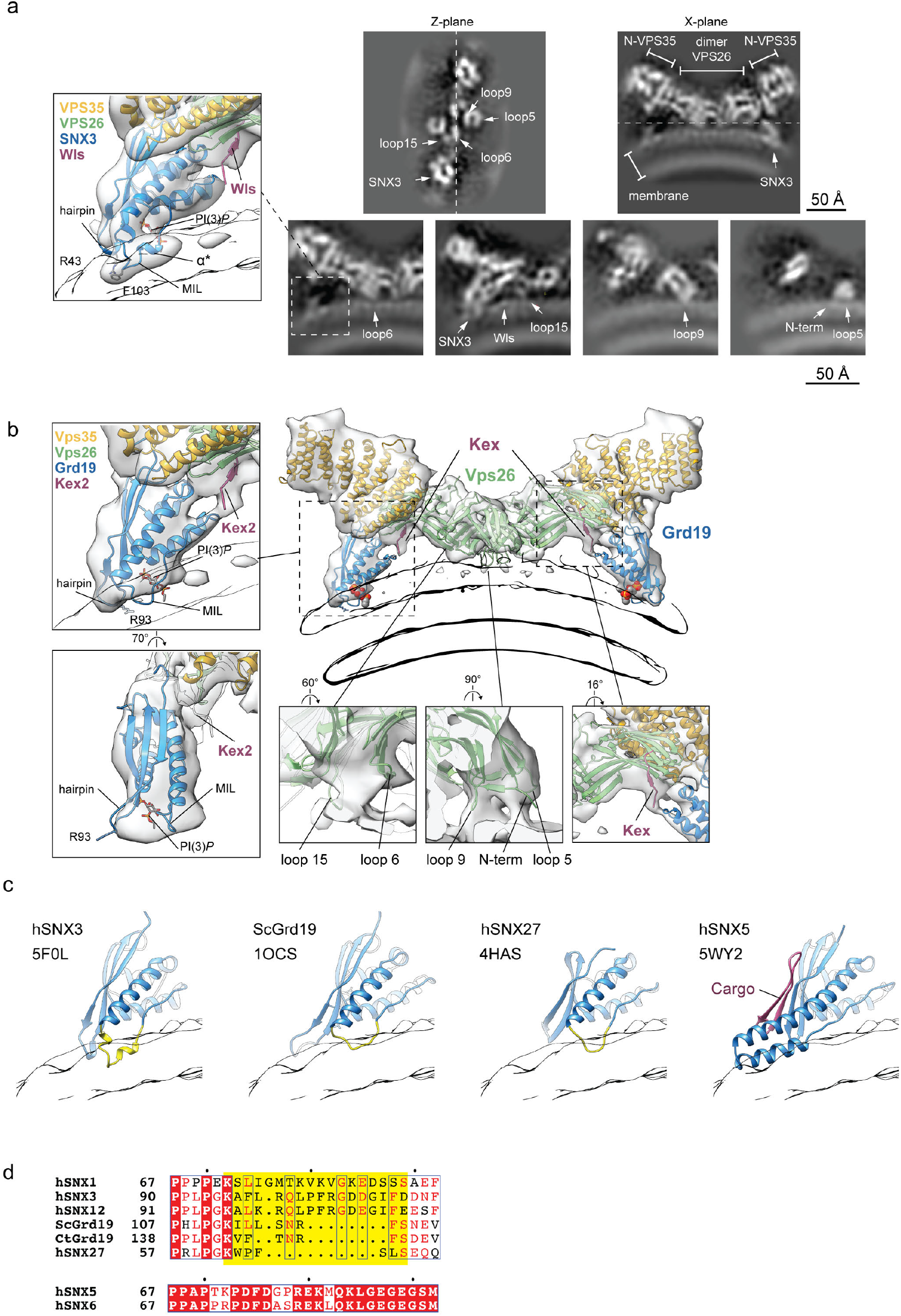
The VPS26:PX region determined by cryo-EM and models of membrane docking of Retromer associated PX domains with different MIL regions. **a,** Top panels: orthoslices through the EM map in the Z plane (view towards the membrane) and X plane (view along the tubule axis) of metazoan retromer:SNX3. Dashed lines show the position of the X-plane in the Z-plane and vice-versa. The membrane, SNX3, VPS26, VPS35, and VPS26 loops are indicated. Bottom panels contain magnified views of X planes through membrane docking elements: SNX3, VPS26 loops, N-terminus of VPS26 and Wls cargo peptide. The inset on the left shows the boxed region as an overlay of semi-transparent EM map and ribbon model demonstrating SNX3 membrane anchoring (as in **Fig2b**). **b**, The fungal retromer:Grd19 coat is shown from the same set of views as metazoan retromer:SNX3 in **Fig2b** to facilitate comparison. **c**, Models of the membrane docking of Retromer-associated PX domains which differ in the MIL region (h-human, Sc – yeast). The position of the PX domain relative to the membrane was modeled by superimposing high resolution crystal structures (PDBs are indicated) with the SNX3 PX domain from the metazoan retromer complex. The membrane is shown as an outline and the MIL region is highlighted in yellow. MIL length and modelled membrane penetration differ between hSNX3, its yeast ortholog ScGrd19 and hSNX27 PX domains. In the hSNX5 PX domain, the MIL is replaced by a helix-turn-helix element that forms a cargo-binding pocket and does not anchor the PX domain in the membrane. **d,** Sequence alignment of MIL regions of PX domains of key retromer-associated SNX proteins including these shown in **c.** Upper panel: PX domains with MIL region (outlined in yellow); Lower panel: PX domains of PX-BARs SNX5 and SNX6 with a helix-turn-helix element.

**Fig S4.**
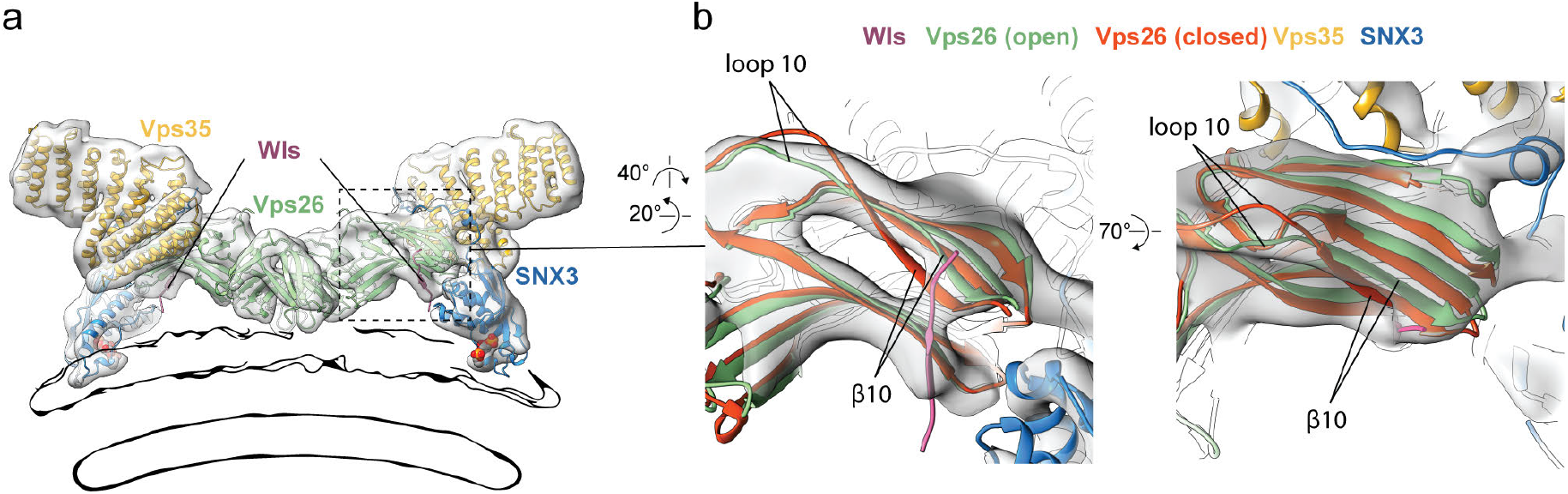
VPS26 is in the open conformation in the membrane-assembled retromer:SNX3 coat. **a.** An overview of the VPS26 dimer region from **Fig2b** with color coded subunits. The boxed region is rotated and magnified in **b. b,** Rigid body fits of VPS26 in its open state (light green, PDB 5F0L^33^) and closed state (orange, PDB 2FAU^34^). The open form of VPS26 fits well into the EM map while for the closed form of VPS26, β10 and the downstream loop10 protrude from the EM map.

**Fig S5.**
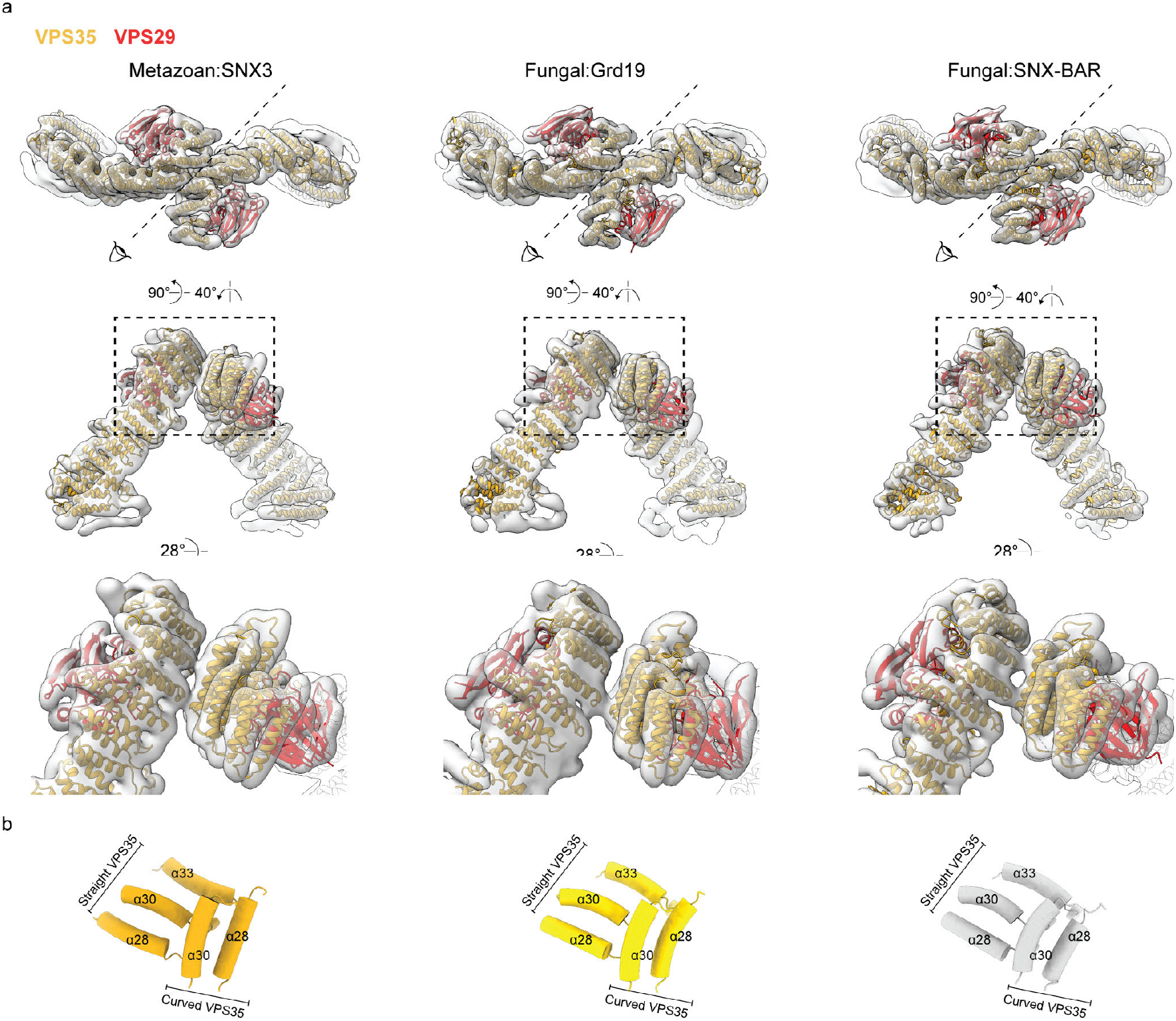
Asymmetric VPS35 arches are a conserved feature of the retromer coat. **a**, VPS35 and VPS29 pseudo-atomic models depicted by ribbons and overlaid with semi-transparent EM maps. **b**, pipe depiction of helices that are involved in VPS35 dimerisation.

**Fig S6.**
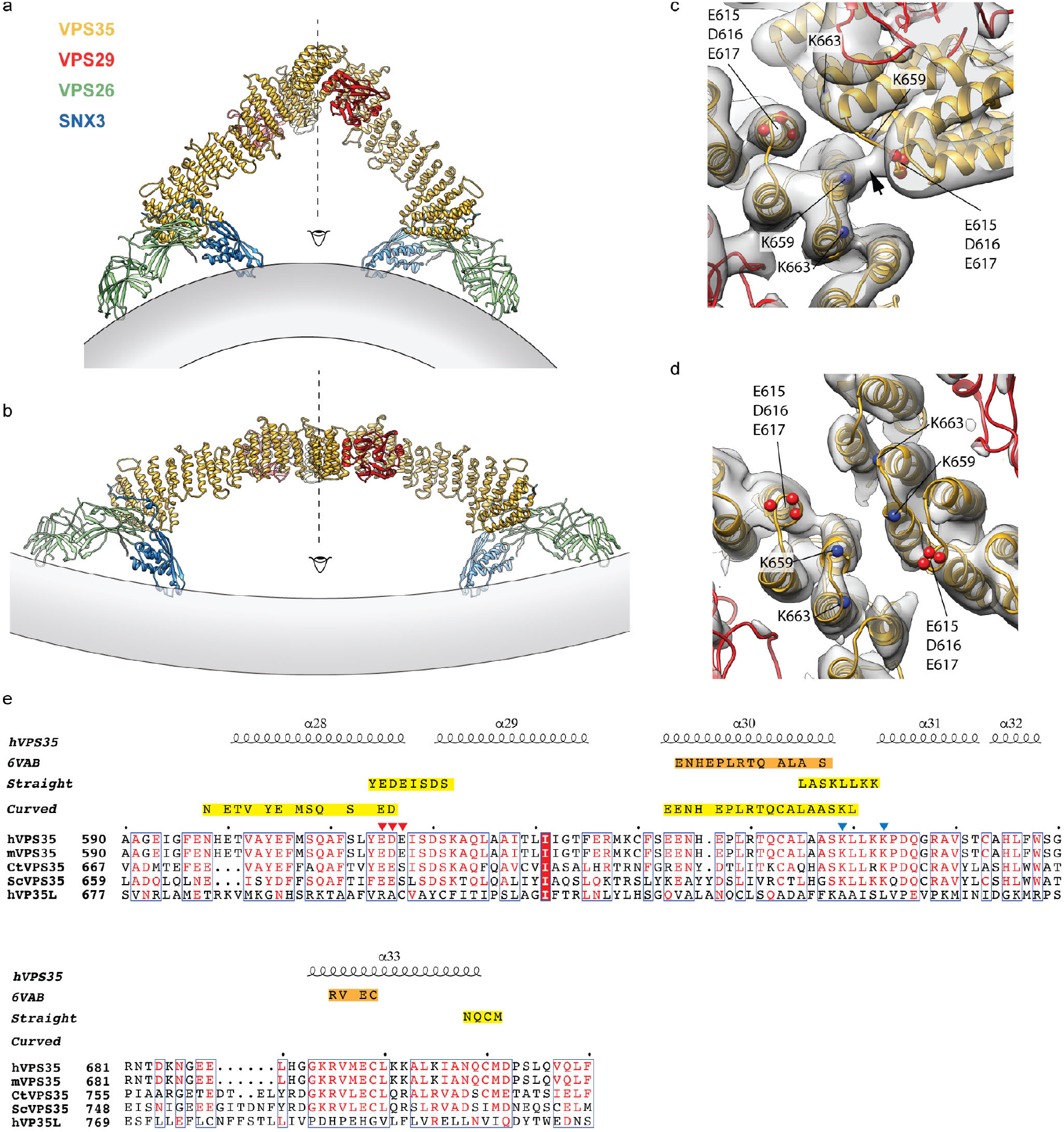
Comparison of metazoan retromer structures determined on the membrane or in solution. **a**, Ribbon models of retromer:SNX3 of membrane-assembled coat determined in this work. The position of the membrane is illustrated. **b,** Ribbon models of retromer:SNX3 based on a structure determined in solution at low ionic strength conditions (70mM) by single particle analysis (PDB 6VAB, EMDB 21135)^21^. In-solution VPS35 dimerisation results in a “flat” assembly different to the arch-like assembly of VPS35 in the membrane-assembled coat. The inverse membrane curvature that would be required in order to bind both VPS26 and SNX3 in this model is illustrated. Kendall et al also presented the solution structure of a retromer arch in which bound SNX3 would be located far from the membrane, and concluded that the arch is unlikely to be relevant to coat assembly. We note that handedness can be difficult to determine in low-resolution single particle structures, and that inverting the handedness of the Kendall et al structure gives an asymmetric arch which is consistent with the asymmetric arch we observed on the membrane, placing SNX3 on the membrane. **c**, Close up view of the VPS35 dimerisation interface observed when retromer is membrane assembled, or **d,** in solution. Cα of charged residues E615, D616, E617 and K659, K653 are marked as red and blue spheres respectively. The acidic E615, D616, E617 cluster in one VPS35 is opposed by the basic cluster K659, K653 in the other VSP35 in the asymmetric interface of membrane-assembled retromer with a clear EM density contact bridge (arrowhead), **c**, while in the symmetrical interface of the in-solution complex these oppositely charged clusters are distant from each other, with positively charged K659s in closest proximity, **d**. **e**, Sequence alignment of the dimerisation regions in VPS35 from different organisms (h- human, m- mouse, Ct-*Chaetomium thermophilum*, Sc - yeast) as well as the VPS35 homologue from the retriever complex (hVPS35L). Secondary structure elements are marked. E615, D616, E617 and K659, K653 are marked as red and blue triangles respectively. Residues participating in the asymmetric VPS35 dimer interface from this study shown in **c**(defined at 10 Å threshold of backbone to backbone distance) are indicated in yellow boxes. The asymmetric interface is formed by residues in helices α28, α30 in the more curved VPS35 and the loops between α28-α29, α30-α31 and α33-α34 in the straighter VPS35. The symmetrical VPS35 dimer interface from **c**(PDB 6VAB) is indicated by the orange box. The symmetrical dimerisation interface results in an inter-subunit contact between α30 helices that does not include any of the mentioned charged residues at the 10 Å threshold.

**Fig S7.**
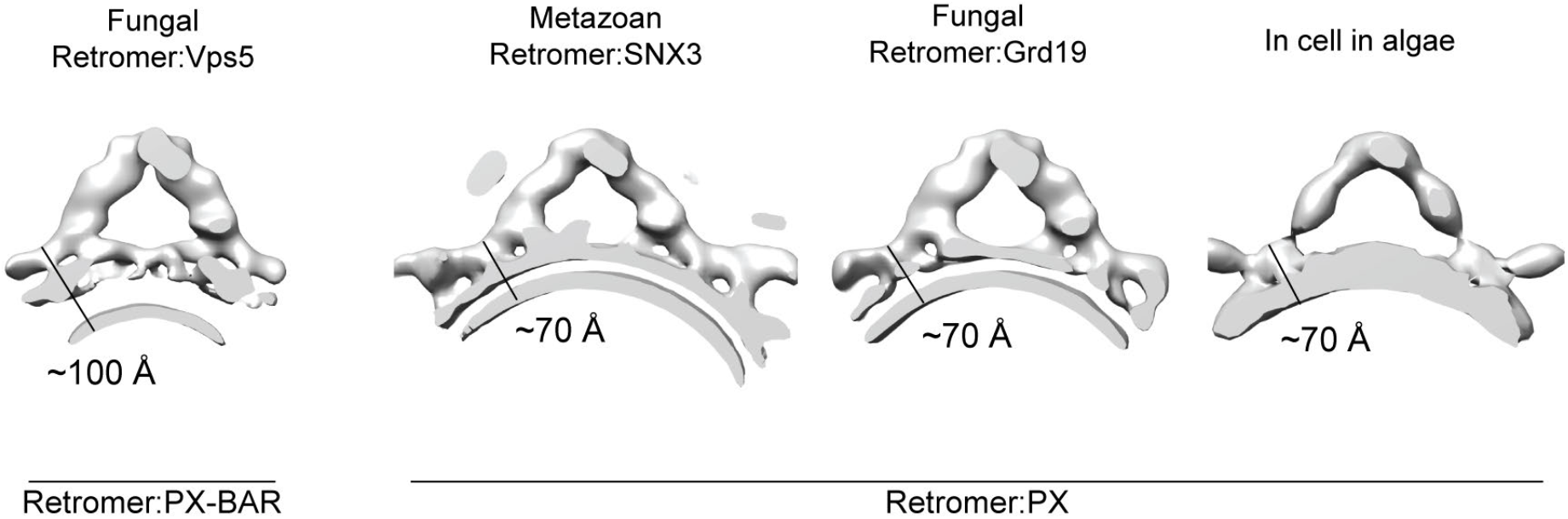
A comparison of available structures of membrane-associated Retromer. EM maps were filtered to 35 Å. In the retromer:Vps5 structure, retromer is bound to the PX-BAR layer, while in metazoan and fungal SNX3(Grd19) complexes, Retromer binds the membrane directly via the VPS26 dimer. The resulting difference in the distance between the retromer arch and the membrane can be distinguished even at low resolution. The approximate distances between the top of the VPS26 dimer and the inner membrane layer in retromer:PX-BAR and retromer:PX complexes correspond to ~100 Å and ~70 Å respectively. These measurements suggest that the majority of retromer arches previously identified in situ within cells of the green algae *Chlamydomonas reinhardtii* (EMDB 0161^6^) are docked directly onto the membrane via VPS26 consistent with a retromer:PX binding mode.

**Table S1.** Cryo-electron tomography data collection.

|  | Retromer:Grd19:Kex2 | Retromer:SNX3:Wls |
| --- | --- | --- |
| Microscope, Voltage (keV) | Titan Krios, 300 |  |
| Detector | Gatan Quantum K2 | Gatan Quantum K3 |
| Energy filter slit width (eV) | 20 |  |
| Electron exposure (e/Å<sup>2</sup>) dose fractionation | ~130, uniformly distributed over tilt series |  |
| Defocus range (μm) | 1.5 – 4.0 | 1.0 -3.5 |
| Tilt scheme (min/max, step) | -60°/+60, 3°, dose-symmetrical (Hagen scheme) |  |
| Movie recording | 10 frames per tilt |  |
| Magnification (times) | X105,000 | X53,000 |
| Pixel size (Å) | 1.379 | 1.701 |
| Number of tomograms acquired/used (no.) | 84/82 | 139/113 |
| Traced membranes (no.) and luminal diameter (mean±std) | 245 tubes (34±2 nm) | 219 tubes (45±8 nm) |

**Table S2.** Subtomogram averaging image processing parameters and statistics.

|  | Retromer:Grd19:Kex2 |  | Retromer:SNX3:WIS |  |
| --- | --- | --- | --- | --- |
|  | arch | Vps26 dimer | arch | VPS26 dimer |
| EMDB/PDB ID | XXXX | XXXX | XXXX | XXXX |
| Initial oversampled subtomograms (no.) | 449,807 |  | 822,112 |  |
| Subtomograms after distance/cluster cleaning (no.) | 55,661 | 52,619 | 112,945 | 97,060 |
| Subtomograms after cross-correlation threshold or edge cleaning (no.) | 46,633 | 49,886 | 89,313 | 66,271 |
| Final subtomograms (no.) | 46,633 | 49,886 | 89,313 | 66,271 |
| Symmetry imposed | none |  |  |  |
| Resolution at FSC threshold 0.143 (Å) | 9.5 | 9.2 | 8.9 | 9.5 |
| Resolution range (Å) | 7-11 | 8-16 | 7-11 | 8-16 |

